# Structure of Orthoreovirus RNA Chaperone σNS, a Component of Viral Replication Factories

**DOI:** 10.1101/2023.07.31.551319

**Authors:** Boyang Zhao, Liya Hu, Soni Kaundal, Neetu Neetu, Christopher H. Lee, Xayathed Somoulay, Banumathi Sankaran, Gwen M. Taylor, Terence S. Dermody, B. V. Venkataram Prasad

## Abstract

The reovirus σNS RNA-binding protein is required for formation of intracellular compartments during viral infection that support viral genome replication and capsid assembly. Despite its functional importance, a mechanistic understanding of σNS is lacking. We conducted structural and biochemical analyses of an R6A mutant of σNS that forms dimers instead of the higher-order oligomers formed by wildtype (WT) σNS. The crystal structure of selenomethionine-substituted σNS-R6A reveals that the mutant protein forms a stable antiparallel dimer, with each subunit having a well-folded central core and a projecting N-terminal arm. The dimers interact with each other by inserting the N-terminal arms into a hydrophobic pocket of the neighboring dimers on either side to form a helical assembly that resembles filaments of WT σNS in complex with RNA observed using cryo-EM. The interior of the crystallographic helical assembly is positively charged and of appropriate diameter to bind RNA. The helical assembly is disrupted by bile acids, which bind to the same hydrophobic pocket as the N-terminal arm, as demonstrated in the crystal structure of σNS-R6A in complex with bile acid. This finding suggests that the N-terminal arm functions in conferring context-dependent oligomeric states of σNS, which is supported by the structure of σNS lacking the N-terminal arm. We discovered that σNS displays RNA helix destabilizing and annealing activities, likely essential for presenting mRNA to the viral RNA-dependent RNA polymerase for genome replication. The RNA chaperone activity is reduced by bile acids and abolished by N-terminal arm deletion, suggesting that the activity requires formation of σNS oligomers. Our studies provide structural and mechanistic insights into the function of σNS in reovirus replication.

## Introduction

Most viruses that replicate in the cytoplasm of host cells form neoorganelles that serve as sites of viral genome replication and particle assembly. These highly specialized viral factory (VF) structures concentrate viral replication proteins and nucleic acids, limit activation of cell-intrinsic defenses, and coordinate release of progeny particles. Both viral and cellular proteins contribute to VF formation and orchestrate viral replication functions.

Mammalian orthoreoviruses (reoviruses) are members of the Orthoreovirus genus in the *Reoviridae* family. Reoviruses are classified into four serotypes based on sequence and antigenicity of the σ1 viral attachment protein and include prototype strains type 1 Lang, type 2 Jones, type 3 Dearing, and type 4 Ndelle ^1^. While reovirus can infect many mammalian species, including humans, symptomatic infection develops only in neonatal mammals and human infants and children ^2^. Infection by some reovirus strains appears to induce the development of celiac disease, a complex immune disorder characterized by an immune response to dietary gluten that can damage the intestinal lining and result in diarrhea and malabsorption ^3^. Reovirus can induce apoptosis, necroptosis, and pyroptosis in tumor cells by activating programmed cell death pathways ^4^. Because of these characteristics, reovirus was one of the first viruses used as an oncolytic agent ^5^.

Reovirus forms nonenveloped particles with an icosahedral capsid, 800 Å in diameter, consisting of two concentric protein layers and distinctive turret-like spikes projecting from the twelve icosahedral vertices. Encapsidated within the innermost protein layer is the viral genome, consisting of ten segments of linear double-stranded (ds) RNA, which encode eleven viral proteins. Eight of these proteins are structural and form the two capsid layers, turret, attachment protein, and an internally located RNA-dependent RNA polymerase. There are three nonstructural proteins that function to promote viral replication. Reovirus replication occurs in the cytoplasm. After binding to cellular receptors, the virus particle enters many types of cells by clathrin-mediated endocytosis ^6^. Within endosomes, proteolysis of outer-capsid proteins yields infectious subvirion particles, which penetrate endosomal membranes to release transcriptionally active viral cores into the cytoplasm ^7^. Following transcription of dsRNA segments within the core interior, capped viral mRNAs are extruded through the turrets into the cytoplasm. The capped transcripts serve as templates for translation as well as synthesis of minus-strand genomic RNA by the viral polymerase to produce the dsRNA genome segments. Assortment of the genome segments into progeny core particles and addition of outer-capsid proteins occur inside VFs. Nonstructural proteins σNS and µNS are required to nucleate these compartments during infection.

In addition to its essential function in formation of viral replication factories, σNS recruits viral RNA ^8^ and both the eukaryotic translation initiation factor 3 subunit A (eIF3A) and the ribosomal subunit pS6R to enhance viral RNA translation ^9^. Its association with the major stress granule effector protein, Ras-GAP SH3-binding protein 1 (G3BP1), disrupts stress granule formation, which relieves a block to reovirus replication ^10^. σNS binds viral ssRNA nonspecifically ^11^, protects ssRNAs from degradation, and ferries viral mRNAs to VFs ^12^. A mechanistic understanding of this multifunctional protein is lacking without its atomic-resolution structure. Our previous studies using cryo-EM show that σNS forms heterogeneous, higher-order oligomers and, when associated with ssRNA, it forms filamentous structures ^12^. However, neither the unliganded nor the RNA-bound form of σNS are conducive to high-resolution structural studies.

We found that an arginine-to-alanine mutation at residue 6 in σNS (σNS-R6A) prevents RNA binding and formation of higher-order oligomers, thus facilitating σNS structural and biochemical characterization. From these analyses, we discovered that domain-swapping interactions of the flexible N-terminal arms of σNS dimers facilitate the association of dimers into filamentous structures that can bind ssRNA. Moreover, we found that such assemblies are required for an RNA chaperone activity of σNS, which we postulate is required for reovirus replication. Bile acids bind to the same hydrophobic pocket as the N-terminal arms and disrupt filament formation and impede RNA chaperone activity. Collectively, these findings provide mechanistic understanding of σNS function in reovirus replication and provide clues about dsRNA synthesis in a *Reoviridae* virus.

## Results

### R6A mutation disrupts σNS RNA binding and oligomerization

Initial attempts to crystalize wildtype (WT) σNS were unsuccessful due to binding of the protein to RNA, which leads to formation of higher-order oligomers that disassemble into heterogeneous populations during purification and crystallization. The N-terminal 38 residues of σNS include several amino acids required for σNS binding to RNA ^12^. These residues are conserved in all reovirus σNS sequences reported to date ^13^. We previously engineered σNS mutants incapable of binding RNA to assess the requirement of σNS RNA-binding activity in reovirus replication ^8^.

The σNS-R6A mutant does not distribute to VFs and fails to transport viral mRNA to these structures ^8^. Based on its lack of RNA-binding capacity, we selected the σNS-R6A mutant for purification and crystallization. Expression and purification of σNS-R6A yielded pure and homogenous recombinant protein. Following anion-exchange chromatography, we compared recombinant WT σNS and σNS-R6A for interactions with nucleic acid (**Fig. 1a**). Anion-exchange chromatography of WT σNS showed two peaks in contrast to that of the σNS-R6A mutant, which showed one peak, consistent with the failure of σNS-R6A to bind RNA. Size-exclusion chromatography of WT σNS following anion-exchange chromatography, yielded an oligomer of a size corresponding to a decamer (**Fig. 1b**) in contrast to σNS-R6A, which showed a peak corresponding to a dimer (∼80 kDa). Surprisingly, the selenomethionine (Se-Met)-substituted σNS-R6A, which we purified for crystallization attempts, showed a size-exclusion chromatography profile suggesting multimeric association with a peak corresponding to an octamer.

**Fig. 1.**
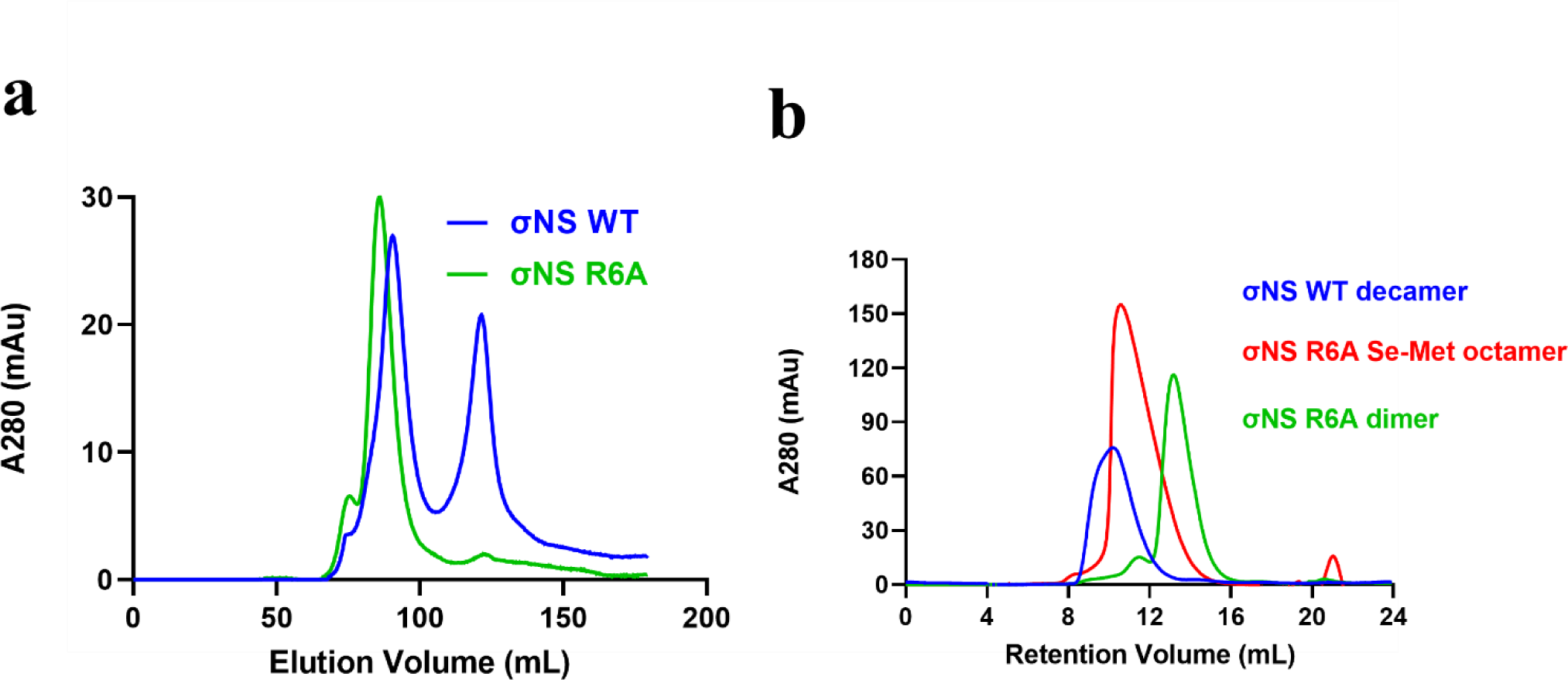
Purification of σNS-R6A dimer. Elution profiles of WT σNS and σNS-R6A after **a** anion-exchange chromatography and **b** size-exclusion chromatography. A size shift is observed with the different elution volumes for each peak.

### σNS forms a stable core structure with a flexible N-terminal arm

To gain a mechanistic understanding of biological functions of σNS, we determined the structure of the σNS-R6A mutant. Although we successfully obtained diffracting crystals of σNS-R6A, determination of the structure by molecular replacement was not possible without suitable phasing models. Accordingly, we proceeded with Se-Met-substituted σNS-R6A and determined the structure using SAD phasing methods to a resolution of 3.0 Å (**Table 1**). Se-Met σNS-R6A crystallized in the space group P6_5_ with two molecules in the asymmetric unit. The crystal structure of σNS-R6A reveals a stable core structure with a protruding N-terminal arm (**Fig. 2a**). The core structure is formed by 13 β-strands and 19 α-helices connected by β- or γ-turns (**Fig. 2b**). The β-strands form four, antiparallel β-sheets on the surface of a globular central body formed by α-helices. The N-terminal residues 4 to 11 form the first helix, which is connected to a short strand by a seven-residue loop. A nine-residue β-hairpin follows the short strand and connects to the first antiparallel sheet of σNS. This antiparallel sheet from residues 30 to 40 together with the N-terminus forms a cleft of about ∼ 25 Å in width. There are several long helices connected by β-turns from residues 245 to 335. The C-terminal region of σNS is connected to helix 19 by a γ-turn and consists of four strands.

**Table 1:** X-ray diffraction data and structure refinement statistics.

| PDB ID | 8TKA | 8TL8 | 8TL1 |
| --- | --- | --- | --- |
| Wavelength (Å) | 0.99 | 0.99 | 0.99 |
| Resolution range (Å) | 34.28 – 3.0 (3.10 - 3.0) | 34.22 - 3.20 (3.31 -3.20) | 34.52-3.16 (3.27 - 3.16) |
| Space group | P 6<sub>5</sub> | P 4<sub>1</sub> | C 1 2 1 |
| Unit cell |  |  |  |
| a, b, c (Å) | 137.11, 137.11, 98.24 | 54.12, 54.12, 305.52 | 77.52, 76.80, 89.09 |
| α, β, γ (°) | 90, 90, 120 | 90, 90, 90 | 90, 94.27, 90 |
| Unique reflections | 41331 (4152) | 11801 (1150) | 7841 (302) |
| Multiplicity | 2.0 (2.0) | 1.0 (1.0) | 1.0 (1.0) |
| Completeness (%) | 99.77 (99.67) | 81.91 (82.38) | 86.39 (33.82) |
| Mean I/σ(I) | 21.07 (3.53) | 18.04 (5.40) | 10.09 (3.21) |
| Wilson B-factor (Å<sup>2</sup>) | 68.54 | 62.51 | 67.74 |
| R-merge (%) | 0.0693 | 0.0311 | 0.1319 |
| R-work (%) | 0.2446 (0.3333) | 0.2240 (0.2944) | 0.208 (0.2561) |
| R-free (%) | 0.2937 (0.4157) | 0.2434 (0.2995) | 0.250 (0.2895) |
| Number of non-hydrogen atoms | 5494 | 5337 | 2708 |
| Macromolecules | 5494 | 5205 | 2637 |
| Ligands | 0 | 132 | 66 |
| Solvent | 0 | 0 | 5 |
| Protein residues | 708 | 669 | 336 |
| R.M.S. deviations |  |  |  |
| Bond lengths (Å) | 0.012 | 0.005 | 0.010 |
| Bond angles (°) | 1.60 | 1.06 | 1.36 |
| Ramachandran |  |  |  |
| Favored (%) | 96.29 | 2.57 | 95.48 |
| Allowed (%) | 3.14 | 2.57 | 3.61 |
| Disallowed (%) | 0.57 | 0.00 | 0.90 |
| Average B-factor (Å<sup>2</sup>) | 81.30 | 59.23 | 67.73 |
| Macromolecules | 81.30 | 58.98 | 67.10 |
| Ligand |  | 69.46 | 95.72 |
| Solvent |  |  | 30.00 |

Numbers in parentheses correspond to the highest resolution range.

**Fig. 2.**
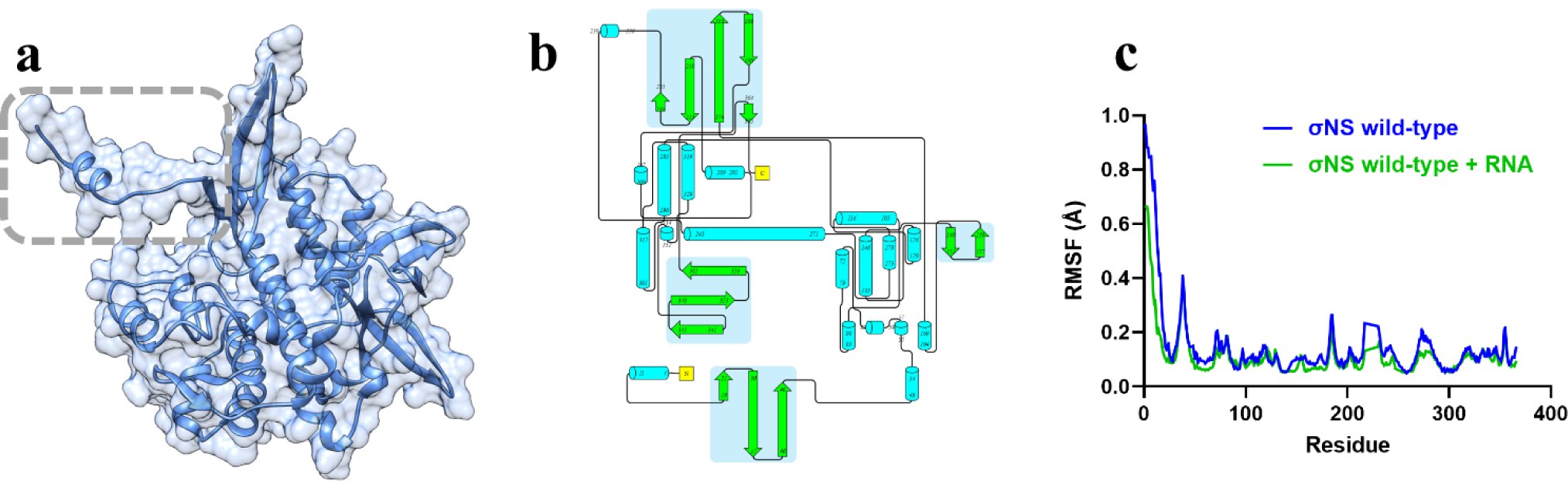
Crystal structure of σNS contains a flexible N-terminal arm. **a** Ribbon and surface representations of the σNS crystal structure. The N-terminal arm is demarcated in a grey-dotted frame. **b** Topologic representation of the σNS crystal structure generated by the PDBsum database. Strands are colored in green with a light blue background, helices are colored in cyan, and the connecting loops are colored in grey. **c** The RMSF graph of the σNS dimer and the σNS dimer in complex with an RNA molecule after 100 nanoseconds of molecular dynamics simulations.

To define flexible regions of σNS, we conducted molecular dynamics simulations based on the σNS-R6A structure. The simulations showed that the σNS N-terminus displays five-fold greater flexibility relative to the other regions of the protein, as assessed by the root mean square flexibility (RMSF) (**Fig. 2c**). The β-sheet connected to the N-terminal loop represents a second region of flexibility, suggesting that σNS contains a flexible N-terminal region. Interestingly, the addition of ssRNA into the simulation diminishes the RMSF of the N-terminus by about 25%.

### σNS forms a stable dimer

The crystallographic asymmetric unit contains two σNS-R6A subunits that form an antiparallel dimer related by non-crystallographic two-fold symmetry, with the N-terminal arms projecting away at either end (**Fig. 3a**). The σNS-R6A dimer is stabilized by an interface with a calculated buried surface area (BSA) of 1853 Å^2^, indicating a biologically relevant oligomer that is stable in solution. The σNS-R6A dimer is stabilized mainly by hydrogen bonds between multiple residues (**Fig. 3b**). Since the dimer interface displays two-fold symmetry, equivalent clusters of residues are formed on both sides of the interface. These clusters include hydrophobic residues (Gly 148 and Val 160) and polar residues (Arg 67, Arg 159, His 164, Asp 168, and Glu 243) (**Fig. 3c**), typical of intersubunit interactions in stable homodimeric proteins ^14^. These observations suggest that σNS dimers with protruding N-terminal arms are the building blocks of higher-order structures.

**Fig. 3.**
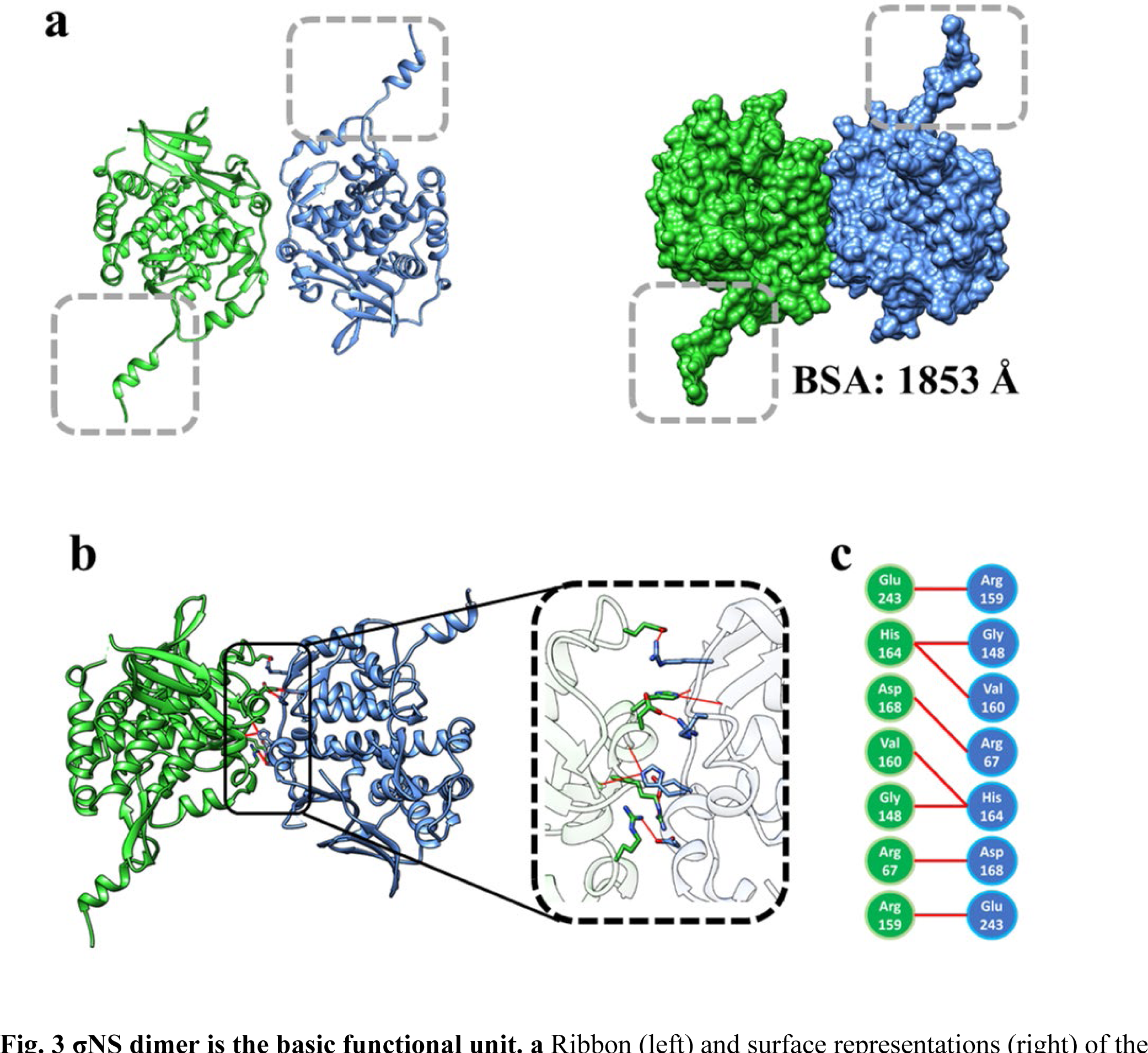
σNS dimer is the basic functional unit. **a** Ribbon (left) and surface representations (right) of the σNS dimer with subunits colored in green and blue. The N-terminal arms are demarcated in grey-dotted frames. **b** Ribbon representation of the σNS dimer highlighting residues that mediate interactions at the dimer interface inside a black frame in blue and green. **c** Amino acid residues from the subunits at the dimer interface are colored blue and green. Hydrogen bond interactions are shown by red lines.

### σNS dimers form a helical assembly by domain-swapping N-terminal arm interactions

The packing of the Se-Met σNS-R6A dimeric subunits in the crystals shows a helical organization with a ∼ 40 Å central cavity formed by domain-swapping interactions between the N-terminal arms of the neighboring dimers (**Fig. 4a-d**). The overall diameter of the helical assembly is ∼ 150 Å. Interactions of σNS to form helical assemblies are consistent with previous observations using cryo-EM, indicating that σNS forms filamentous structures in the presence of RNA ^12^. Such helical assemblies were not expected for the σNS-R6A mutant, as it does not bind RNA and forms only dimers in solution. However, the size-exclusion chromatography profile of Se-Met-substituted σNS-R6A clearly showed that it forms higher-order oligomers in solution as observed with WT σNS (**Fig. 1b**). It is possible that increased hydrophobicity due to substitution of a selenium atom for a sulfur atom^15^ in Met 1 coupled with crystal-packing forces restrict the flexibility of the N-terminal arms and allow this region of the protein to stabilize a multimeric association.

**Fig. 4.**
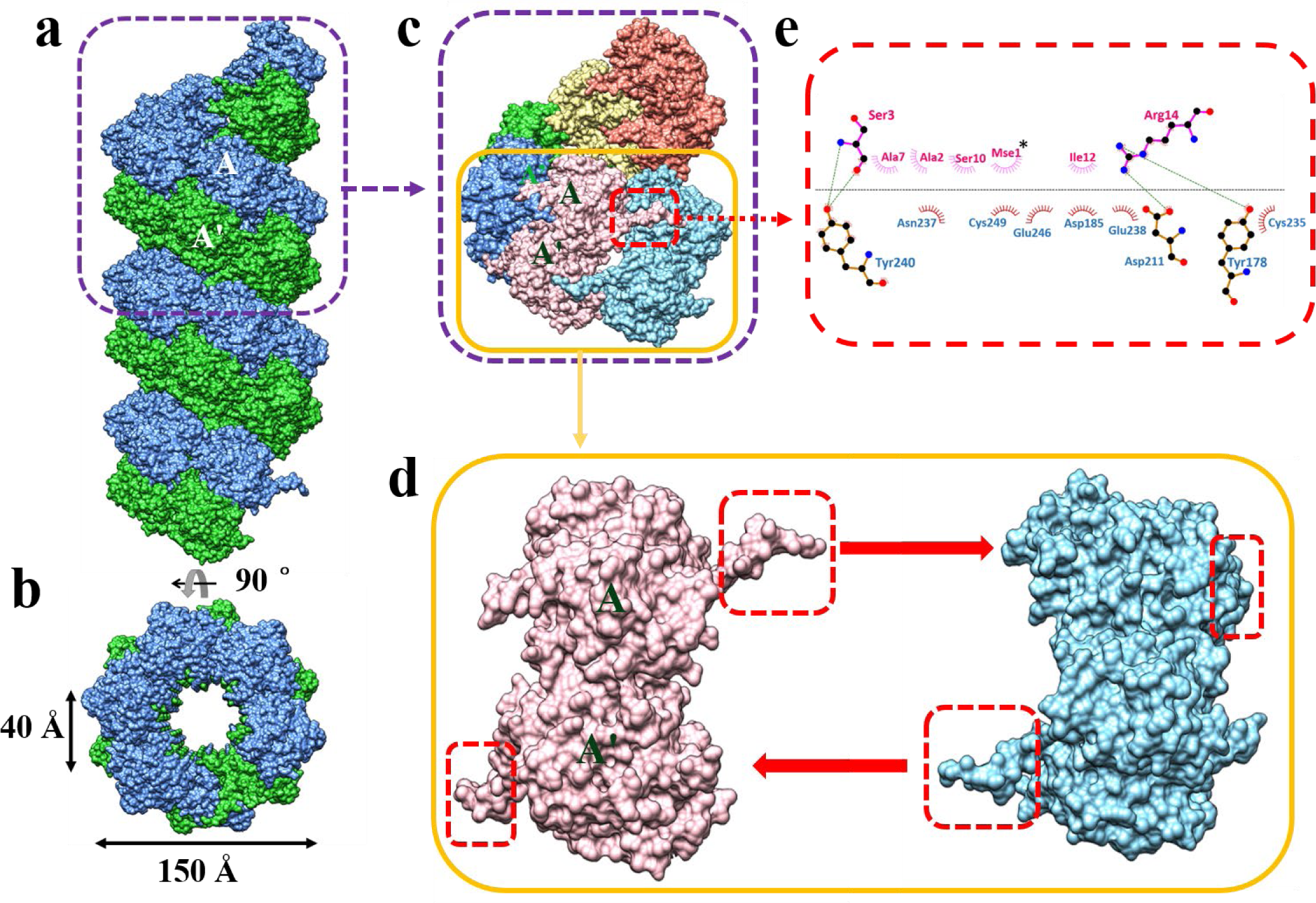
Formation of σNS filaments. **a** Surface representation of the helical assembly formed by σNS dimers using crystallographic P6_5_ symmetry. Dimers are depicted as blue and green subunits. A single helical turn (80 Å in length) is demarcated in the purple-dotted frame. **b** A 90° rotation around the horizontal axis perpendicular to the filament shows the central tunnel. The tunnel is 40 Å in width, and the total diameter of the filament is 150 Å. **c** Single helical turn formed by six σNS dimers. Dimers are depicted in different colors. One pair of interacting dimers (pink and cyan) are demarcated in a yellow frame. The location of the N-terminal arm of the pink dimer is shown within the red frame, with a red arrow pointing to the (**e**) Ligplot of interactions between N-terminal arm residues of the pink dimer and core domain residues of the blue dimer. Residues involved in hydrophobic interactions are shown by brick-red spoked arcs, and hydrogen bond interactions between the residues are indicated by green lines, with carbon, oxygen, and nitrogen atoms depicted in black, red, and blue, respectively. Note the involvement of the Se-Met residue at position 1 (Mse 1), indicated by an asterisk. **d** Interacting dimers (pink and blue) corresponding to the yellow frame in **c** are shown separately to illustrate how the N-terminal arms (in red frame) project away in opposite directions (thick red arrows) to chain-link the neighboring dimers to form the helical assembly shown in Fig. 1a.

### Bile acid derivatives disrupt σNS filament formation

We observed that the native σNS-R6A crystals, without Se-Met substitution, crystallized in the P4_1_ space group instead of P6_5._ Interestingly, native σNS-R6A crystallized only in conditions with bile acid derivatives in contrast to Se-Met σNS-R6A, which crystallized without these additives. Structure determination of native σNS-R6A, using the Se-Met σNS-R6A structure as a molecular replacement model, showed that native σNS-R6A also forms an antiparallel dimer, with two bile acid moieties bound to each subunit in a groove located near the σNS C-terminus (**Fig. 5a**, sites 1 and 2). Binding of bile acid molecules is mediated by hydrophobic contacts with residues Thr 107, Glu 110, Leu 111, Ser 114, Gly 203, Leu 204, Tyr 240, Glu 242, Ala 245, and Glu 246 (**Fig. 5b**). Remarkably, in contrast to the Se-Met σNS-R6A structure, the N-terminal arm is disordered and not observed in the native σNS-R6A structure. Instead of the distinct helical assembly in the structure of Se-Met σNS-R6A, native σNS-R6A dimers crystallized with bile acid show sheet-like packing (**Fig. 5c).** Further examination of the structure showed that one of the bile acid molecules strikingly occupies the same location in the core domain as the N-terminal arm required to form helical assemblies, suggesting that bile acid competes with the N-terminal arm to disrupt the helical assembly formation observed in the Se-Met σNS-R6A structure (**Fig. 5d-f**).

**Fig. 5.**
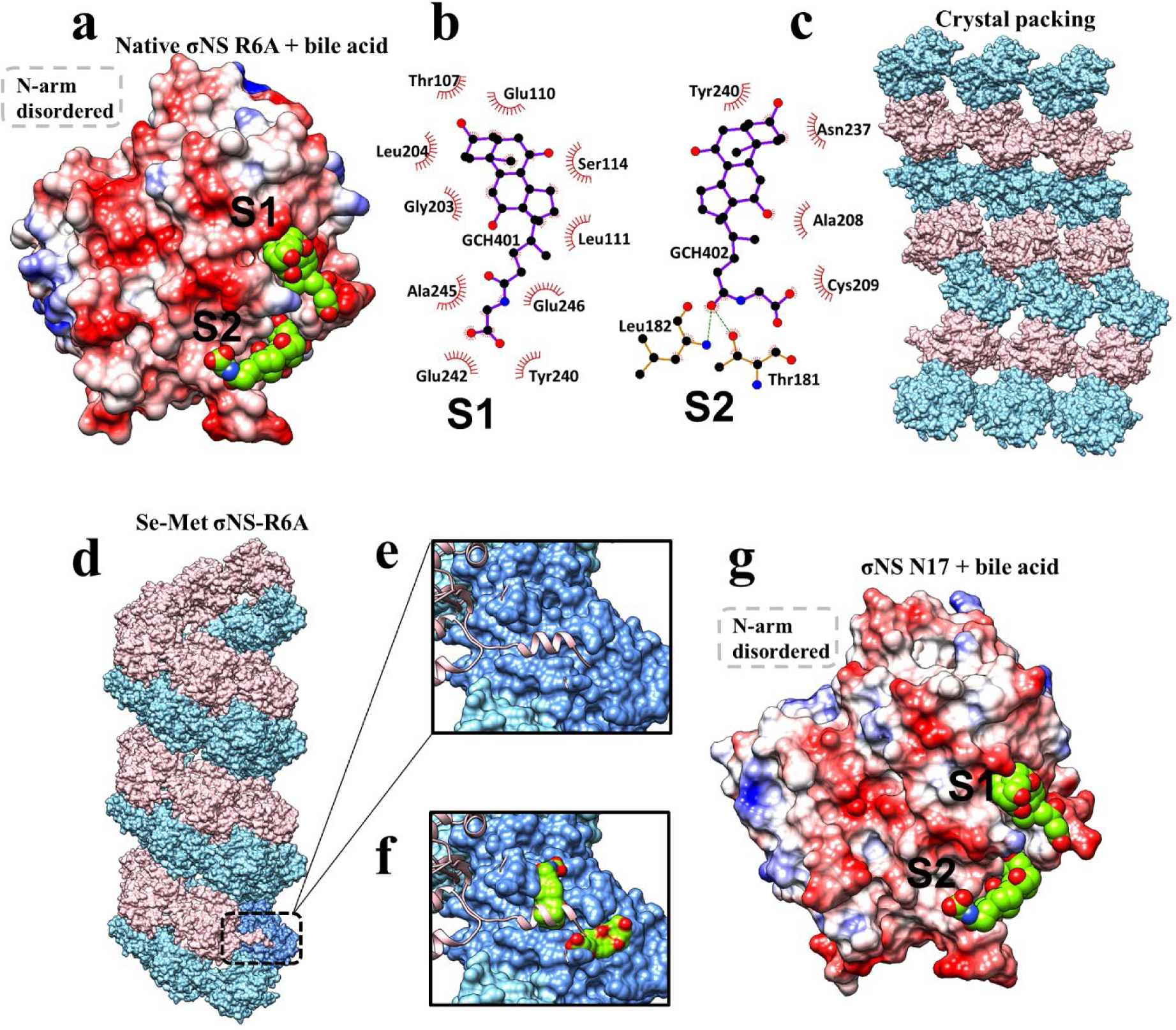
Binding of bile acid derivatives disrupts σNS oligomerization and RNA-destabilizing activity. **a** Electrostatic representation of σNS-R6A in complex with bile acid derivatives (S1 and S2), colored in green. The grey-dotted box demarcates the missing disordered N-terminal arm. **b, c** Residues mediating bile-acid-derivative binding at S1 and S2 defined by LigPlot following the same notations as in Fig. 4e for hydrophobic and hydrogen bond interactions. Bile acid derivatives are shown in purple bonds with carbon and oxygen atoms in black and red, respectively. **c** The higher-order structure formed by crystal packing of σNS-R6A when bile acid derivatives are present. **d** The helical assembly as observed in the crystal structure of Se-Met σNS-R6A with dimeric subunits shown in pink and blue. **e** Close-up view of the N-terminal arm (pink) interacting with a groove in the neighboring subunit (blue). **f** In the structure of σNS-R6A in complex with bile acid derivatives, the same groove is occupied by a bile acid replacing the N-terminal arm, which becomes disordered. **g** Electrostatic representations σNS-ΔN17 in complex with two bile acid derivatives (in green) bound at the same location as in the σNS-R6A structure shown in Fig. 5a. The grey-dotted box demarcates the missing N-terminal arm.

### The σNS N-terminal arm is required for multimeric association

To provide additional evidence for the role of domain-swapping interactions of the N-terminal arms in forming oligomeric assemblies, we engineered a deletion construct, σNS-ΔN17, by truncating the N-terminal 17 residues of the protein. The σNS-ΔN17 mutant formed dimers in solution and, interestingly, it also crystallized in the presence of bile acid derivatives. The σNS-ΔN17 crystal structure determined using molecular replacement showed a dimer formed by a central core identical to that in the structures of Se-Met or native σNS-R6A (**Fig. 5g**) with two bound bile acid moieties. As expected, no helical packing was observed in the crystal structure. Therefore, the structure of σNS-ΔN17 suggests that the N-terminal arm and the core of WT σNS operate as independent structural units, and removing the N-terminal arm does not alter the dimeric association of the core. Collectively, the three crystal structures provide strong evidence that (i) WT σNS has a tendency to form dimers, (ii) the dimer is the basic unit of multimeric assembly, and (iii) domain-swapping interactions of the flexible N-terminal arms are required to link the dimeric units to form a multimer. Furthermore, the σNS-ΔN17 structure showed that bile acid derivatives bind to the same location in σNS as the N-terminal arms in the σNS-R6A structure (**Fig. 5g**), suggesting that this location in the σNS core domain has intrinsic affinity for cholesterol-like molecules and that binding to such molecules disrupts filamentous helical assemblies by competing with the binding of the N-terminal arms.

### σNS oligomer exhibits RNA chaperone activity

The surface representation of the helical assembly in the Se-Met σNS-R6A structure shows a negatively charged exterior (**Fig. 6a**) and a positively charged interior (**Fig. 6b**). A cutaway view of the central tunnel of the helical assembly reveals a substantial number of positively charged residues, suggesting that this region can bind RNA by electrostatic interactions. *In silico* docking experiments demonstrated that the central tunnel can accommodate an RNA strand surrounded by σNS dimers (**Fig. 6c-e**). This finding, coupled with previous studies showing that avian reovirus σNS displays RNA-chaperone activity^16^, prompted us to hypothesize that mammalian orthoreovirus σNS is likewise an RNA chaperone. To test this hypothesis, we first examined whether purified WT σNS and σNS-R6A disrupt dsRNA by incubating each of these σNS proteins with an RNA-RNA duplex with an overhang of 6 nucleotides on either side. After incubation, the reaction mixtures were treated with proteinase K and resolved by electrophoresis in nondenaturing polyacrylamide gels. The RNA-RNA duplex migrated as a single prominent band when σNS was not added to the reaction mixtures (**Fig. 6f**). However, following addition of increasing concentrations of WT σNS, we observed a proportional increase in the formation of ssRNA molecules. As an important control, formation of ssRNA was not observed in a similar experiment using σNS-R6A (**Fig. 6g**). These results indicate that σNS catalyzes formation of ssRNA molecules from dsRNA and that σNS displays RNA-destabilizing activity. We next evaluated whether σNS is capable of annealing two ssRNA molecules to form dsRNA. In this experiment, we observed a proportional increase in the formation of dsRNA molecules following addition of increasing concentrations of WT σNS to the reaction mixtures (**Fig. 6h**), indicating that σNS catalyzed formation of dsRNA molecules from ssRNA and thus possesses RNA-annealing activity. Since the reaction buffer for these experiments contained no ATP, the RNA chaperone activity of σNS is independent of energy input. These experiments demonstrate that σNS is not a helicase but an RNA chaperone.

**Fig. 6.**
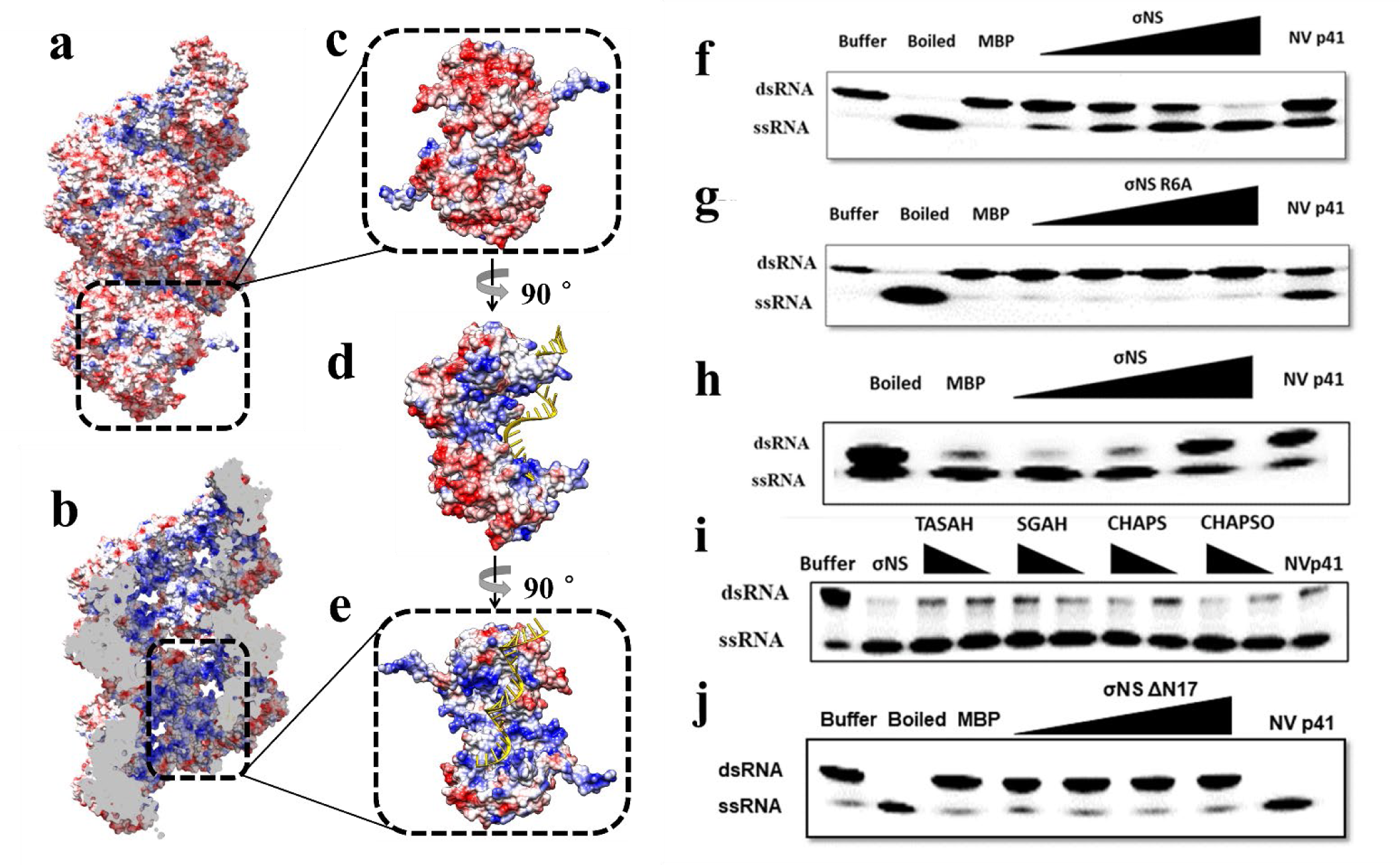
RNA chaperone activity of σNS. **a** Electrostatic representation of a σNS filament with red indicating negatively charged residues and blue indicating positively charged residues. The exterior of the filament is negatively charged. **b** Cutaway view of the filament. The interior of the filament is positively charged. **c** A single dimer is demarcated in a black-dotted frame with its relative position within the filament. **d** A 90-degree rotation around the z-axis from the previous position shows a side-view of the dimer. A docked RNA molecule is colored in yellow. **e** A 90-degree rotation around the z-axis from the previous position shows the positively charged pocket of σNS that binds to a docked RNA molecule colored in yellow. The black-dotted frame depicts the position of the dimer within the filament. **f** RNA-helix destabilizing activity of WT σNS. Boiling of the duplex disrupts duplex formation. Increasing concentrations of WT σNS yield more ssRNA products. Maltose-binding protein (MBP) and norovirus (NV) P41 serve as negative and positive controls, respectively. **g** RNA-helix destabilizing activity of σNS-R6A. Increasing concentrations of σNS-R6A yield only limited ssRNA products. **h** RNA-annealing activity of WT σNS. Boiling two ssRNA molecules promotes strand annealing. Similar levels of annealing activity are observed following incubation with the highest concentration of WT σNS. **i** Effects of each component of the bile acid derivatives on the RNA-destabilizing activity of σNS. TASAH, taurocholic acid sodium salt hydrate. SGAH, sodium glycocholate hydrate. CHAPS, 3-[(3-cholamidopropyl)dimethylammonio]-1-propanesulfonate. CHAPSO, 3-[(3-cholamidopropyl)dimethylammonio]-2-hydroxy-1-propanesulfonate. **j** RNA-helix destabilizing activity of σNSΔ17. Increasing concentrations of σNS Δ17 does not yield significant ssRNA products.

### Binding of bile acid inhibits σNS chaperone activity

From our observation that bile acid binding disrupts formation of filamentous helical assemblies of σNS, we hypothesized that bile acid binding would impede σNS RNA chaperone activity. To test this hypothesis, we conducted RNA chaperone assays following incubation of σNS with different components of bile acid derivatives, including taurocholic acid sodium salt hydrate (TASAH), sodium glycocholate hydrate (SGAH), 3-[(3-cholamidopropyl)dimethylammonio]-1-propanesulfonate (CHAPS), and 3-[(3-cholamidopropyl)dimethylammonio]-2-hydroxy-1-propanesulfonate (CHAPSO). Each of these components reduced the RNA-destabilizing activity of σNS to varying degrees (**Fig. 6h**), suggesting that the RNA chaperone activity of σNS requires formation of filament-like assemblies. As anticipated, σNS RNA-binding capacity and oligomer formation were abolished by removal of the N-terminal 17 residues and, concomitantly, the σNS-ΔN17 mutant did not display RNA chaperone activity (**Fig. 6i**).

## Discussion

VFs serve an essential function in the replication of many viruses, as these structures house viral genome synthesis and assembly of progeny virions. In the case of mammalian orthoreoviruses, two nonstructural proteins, σNS and µNS, initiate VF formation, and σNS recruits viral RNAs into these compartments. Our crystallographic and biochemical studies reported here provide a mechanistic understanding of how σNS forms filamentous assemblies by linking dimeric units through domain-swapping interactions of the N-terminal arms and further how such a multimeric association is required for the RNA-helix destabilizing and annealing activities of σNS. These findings implicate σNS in unwinding viral RNA for loading into the viral RNA-dependent RNA polymerase during genome replication and promoting interactions between the viral RNAs to facilitate their selective encapsidation during virion assembly.

### Higher-order oligomers and a flexible element–a common theme in VF formation?

Although distinct in its structure, σNS displays several functional and mechanistic similarities with other dsRNA virus nonstructural proteins, such as rotavirus NSP2 and rice black-streaked dwarf virus (RBSDV) P9-1. Like σNS, both NSP2 and P9-1 are implicated in formation of VFs. The striking common structural feature of these proteins is the formation of oligomeric assemblies requiring a flexible structural element. However, they differ in detail. Unlike the formation of filamentous assemblies by σNS, rotavirus NSP2 forms a donut-shaped octamer ^17,18^. Instead of the flexible N-terminal arm of σNS, which is responsible for dimer-dimer association in filamentous assembly formation, NSP2 has a flexible C-terminal arm, which facilitates inter-octamer association through domain-swapping interactions with neighboring NSP2 molecules ^17^. Formation of higher-order NSP2 octamers is thought to be the basis for VF formation during rotavirus replication ^17,19^. RBSDV P9-1 also forms an octamer using a flexible C-terminal arm^20–22^. Like rotavirus NSP2, higher-order P9-1 octamers are required for formation of VFs, as deletion of this sequence disrupts RBSDV VFs ^22^. From our studies, deletion of the N-terminal arm of σNS abrogates dimer-dimer interactions and precludes formation of higher-order multimers. However, further experiments are required to assess the function of the flexible N-terminal arm in the formation of functional VFs.

### RNA chaperone activity–a flexible N-terminal arm

An important finding from our studies is that mammalian orthoreovirus σNS exhibits helix-destabilizing and RNA-annealing activities without requiring a metal ion or ATP, which makes this protein a *bona fide* RNA chaperone ^24^. Similar RNA chaperone activity has been reported for rotavirus NSP2 ^25^ and avian reovirus σNS ^16^. The RNA duplex used in our experiments contains overhangs at the ends of both strands, suggesting that passive strand displacement activity is initiated after σNS binds to unduplexed overhang regions. The binding of σNS would then prevent reannealing of the duplex and further separate the two strands. Using complementary single-stranded RNAs, our studies show that σNS binds, unfolds, and releases these molecules to facilitate formation of duplex RNA.

The RNA chaperone activity of σNS requires the flexible N-terminal arm with an arginine at residue 6 to function in domain-swapping interactions, as the R6A substitution or deletion of the N-terminal arm curtails this activity. In an analogous scenario, deletion of the flexible C-terminal arm of NSP2 also significantly reduces RNA chaperone activity ^26^. Thus, the requirement for a flexible region in these proteins suggests that the mechanism is likely mediated by disorder or entropy transfer. One possibility is that σNS uses the flexibility of the N-terminal arm for binding RNA and subsequently releases the bound RNA. Such a possibility is consistent with our observation that a single residue, Arg 6, in the N-terminal arm influences the RNA-binding capacity of σNS ^8^. The free N-terminal arms of the σNS multimer subunits (**Fig. 5e**) likely exist in a dynamic equilibrium between a freely exposed open state, in which Arg 6 can initiate binding to RNA, and a closed state, in which the N-terminal arm is engaged in domain-swapping interactions for RNA release. Such dynamics also may allow the N-terminal arm to recruit free σNS dimers to elongate the multimer and coat the RNA as a mechanism for unwinding (**Fig. 7**). The dynamics and preference between open and closed states are likely influenced by the presence of RNA and buffer conditions (pH and ionic strength). This idea is consistent with our observation that in solution σNS exists as an octamer in the absence of RNA, but in the presence of RNA, as observed in our previous cryo-EM images ^12^, σNS forms filamentous structures of varying lengths.

**Fig. 7.**
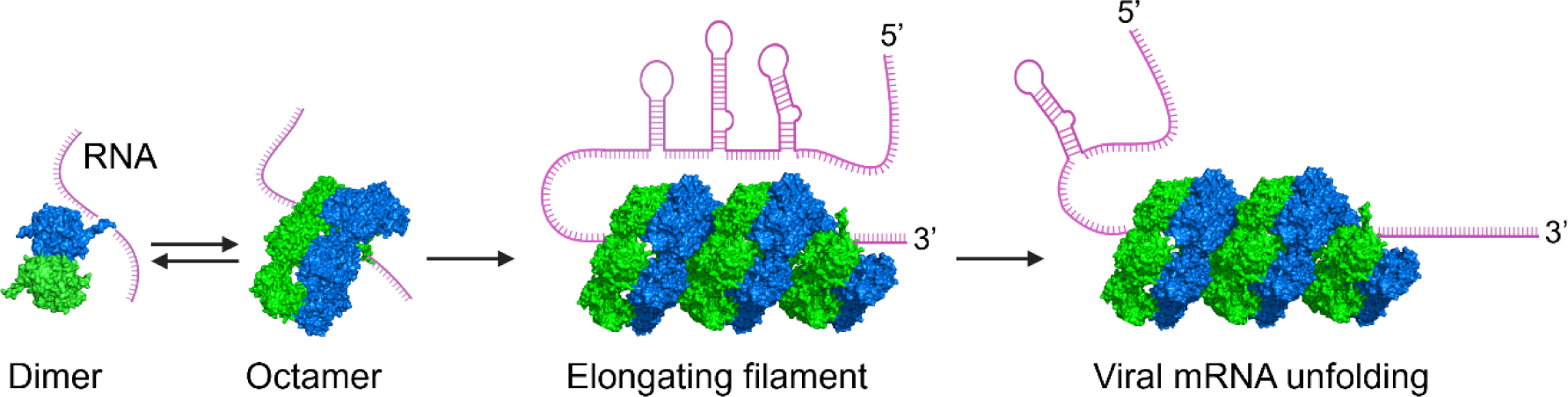
Model of σNS oligomerization, RNA binding, and RNA unfolding. The free N-terminal arms of σNS oligomers exist in equilibrium between an open state that can initiate RNA binding and a closed state that engages in domain-swapping interactions and RNA release from the N-terminal arm. The dynamics of the N-terminal arms allow for recruitment of free σNS dimers to elongating σNS oligomers to coat the RNA and enhance RNA unfolding.

### Bile acid binding–tampering with the flexibility of the N-terminal arm

A serendipitous but fascinating finding from our study is that bile acid derivatives bind to the same site in σNS as the N-terminal arm and impede the RNA chaperone activity of σNS (**Fig. 5h**). These findings are consistent with the proposed mechanism of σNS RNA chaperone activity. First, the σNS crystal structure shows that the free N-terminal arm displaced by bile acid is disordered, as the electron density for this region is absent in the presence of bile acid. This observation is consistent with the disorder or entropy-transfer model of RNA chaperone activity ^27^, in which entropy is transferred to bound RNA to facilitate RNA refolding. Second, our finding that bile acid binding does not entirely abrogate RNA chaperone activity is consistent with the idea that RNA modulates the kinetics of open and closed states of the N-terminal σNS arm. It is possible that differences in the efficiency by which bile salts diminish RNA chaperone activity are determined by differences in the capacity of these molecules to compete with the N-terminal arm in the presence of RNA.

### Role of RNA chaperone activity in reovirus replication

Our discovery that σNS displays RNA chaperone activity raises an important question about the stage in viral replication at which this activity is required. While the function of σNS RNA-binding activity in the recruitment of viral RNAs to VFs has been established ^8^, further studies are needed to unequivocally determine the precise function of σNS RNA chaperone activity. As proposed for RNA chaperones of single-stranded RNA viruses ^27^, σNS RNA chaperone activity may be necessary for unfolding viral mRNA for translation, replication, packaging, or some combination of these steps. In the case of dsRNA viruses like reoviruses and rotaviruses that package multiple dsRNA segments, RNA chaperone activity is likely essential for proper refolding of the kinetically trapped intermediate structures in capped viral mRNAs to allow these RNAs to serve as templates for dsRNA synthesis by the viral polymerase and also for facilitating interactions between specific viral RNAs for gene segment assortment and packaging.

## Methods

### Expression and purification of σNS constructs

Synthesized genes for WT σNS, σNS-R6A, and σNS-ΔN17 were subcloned into the bacterial expression vector pET28 with an N-terminal His tag and a TEV protease cleavage site (Epoch Life Science). *Escherichia coli* DE3 cells (Novagen) were transformed with the pET28 plasmid and induced with 0.5 mM isopropyl β-D-1-thiogalactopyranoside (IPTG) (Millipore Sigma) when the optical density at 600 nm reached 0.6. *E. coli* B834 cells were incubated in Dream medium supplemented to contain Dream Nutrient Mix and 10 mg/mL Se-Met. Cells were resuspended in 50 mM Tris-HCl (pH 8.0), 300 mM NaCl, and 10 mM imidazole and supplemented to contain a protease inhibitor cocktail (Roche). Cells were lysed using a microfluidizer (Microfluidics), followed by removal of cell debris by centrifugation at 39,000 x *g* at 4°C for 30 minutes. His-tagged σNS was loaded onto a Ni-NTA agarose column (Qiagen) and eluted using a gradient of 50 mM Tris-HCl (pH 8.0), 300 mM NaCl, and 250 mM imidazole. The eluted His-tagged σNS was concentrated using a 10-kDa centrifugal filter unit (Millipore), dialyzed into 50 mM Tris-HCl (pH 8.0), 300 mM NaCl, and 10 mM imidazole, and incubated with TEV protease at 4°C overnight. The cleaved protein mixture was reloaded onto a Ni-NTA agarose column to remove the His-TEV protease and uncleaved His-σNS. Cleaved σNS was subjected to ion-exchange chromatography using a Q-column (GE Healthcare) and eluted with 20 mM Tris-HCl (pH 8.0) and 1 M NaCl. Eluted σNS was separated using an S200 16/60-increase column in 10 mM HEPES (pH 8.0) and 150 mM NaCl.

### Crystallization, data processing, structure determination, and refinement

Freshly purified Se-Met σNS-R6A, native σNS-R6A, and σNS-ΔN17 were used for initial crystallization screens immediately following size-exclusion chromatography. In each case, fractions with σNS proteins were collected and concentrated to ∼ 10 mg/mL in buffer containing 10 mM HEPES (pH 8.0) and 150 mM NaCl for crystallization. Crystals were propagated at 20°C by hanging-drop vapor diffusion using a Mosquito crystallization robot (TTP LabTech) and imaged using a Rock Imager (Formulatrix). Each drop contained 0.2 μL of the σNS protein and 0.2 μL of crystallization buffer. Se-Met σNS-R6A produced crystals in the condition containing 0.1 M MES monohydrate (pH 6.5) and 1.6 M magnesium sulfate heptahydrate (Hampton Research), whereas native σNS-R6A and σNS-ΔN17 crystallized in the condition with 1.2 % cholic acid derivatives mix, 0.1 M Buffer System 3 (pH 8.5), and 30 % Precipitant Mix 3 (Morpheus III, Molecular Dimensions). Crystals were transferred into a cryoprotectant solution with 20% glycerol and flash-frozen in liquid nitrogen. X-ray diffraction data for σNS were collected on beamline 5.0.1 at the Advanced Light Source at the Lawrence Berkeley National Laboratory. Diffraction data were processed using the CCP4 software suite ^28^. For Se-Met σNS-R6A, phases and initial electron density maps were calculated by single-wavelength anomalous diffraction (SAD) phasing using SHELX ^29^, and automated model building with experimental phases was conducted using ARP/wARP ^30^ before iterative cycles of manual model building and refinement using COOT^31^ and PHENIX ^32^. Further model building was accomplished using COOT based on the difference maps ^31^ (see validation report for PDB id: 8TKA). The structures of native σNS-R6A and σNS-ΔN17 were determined using molecular replacement with PHENIX ^32^. The bile acid binding in these structures was validated by omit maps (see validation reports for PDB id: 8TL8 and 8TL1). Data collection and refinement statistics following the final refinement cycle are provided in Supplementary Table 1. Interactions between σNS and colic acid derivatives were analyzed using LigPlot+ v.2.1 ^33^. Figures were prepared using Chimera ^34^.

### RNA helix destabilizing assay

RNA helix destabilizing assays were conducted as described ^35^ with modifications. Various concentrations of σNS, σNS-R6A, and σNS-ΔN17 (2.5 to 20 µM) and 0.5 μM dsRNA substrate were incubated at 37°C for 1 h with 25 mM HEPES-KOH (pH 8.0), 100 mM NaCl, 2 mM MgCl_2_, 2 mM DTT, and 5 U RNase-OUT in a reaction volume of 20 μL. Reactions were terminated by the addition of 5 U proteinase K for 15 min and 2.5 μL 5 × loading buffer (100 mM Tris–HCl [pH 7.5], 50% glycerol, and bromophenol blue) and electrophoresed in 4-20% native polyacrylamide gels. Gels were scanned using a Typhoon 9200 imager (GE Healthcare). The effect of bile acids on σNS RNA helix destabilizing activity was determined by incubating 20 and 40 μM of different components of bile acid derivatives, including TASAH, SGAH, CHAPS, and CHAPSO, with 20 μM σNS for 1 h prior to initiation of the helix destabilization assay.

### RNA strand hybridization assay

Various concentrations of σNS (2.5 to 20 µM) were incubated with HEX-labeled and unlabeled RNA strands (0.5 μM each strand) at 37°C for 1 h in 50 mM HEPES-KOH (pH 8.0), 2.5 mM MgCl_2_, 2 mM DTT, 0.01% BSA, and 5 U RNase-OUT in a reaction volume of 20 μl. Reactions were terminated as described for RNA helix destabilizing assays. Two complementary stem-loop-structured RNA strands were used for hybridization assays ^35^. Reactions were resolved in 4-20% native polyacrylamide gels and scanned using a Typhoon 9200 imager.

## Acknowledgements

We are grateful to members of the Dermody and Prasad laboratories for many useful discussions. This work was supported by U.S. Public Health Service award R01 AI032539, the Heinz Endowments (T.S.D.), and grant Q1279 from Robert Welch Foundation (B.V.V.P). We acknowledge the Advanced Light Source (8.2.2) (Berkeley, CA) for X-ray data collection. The ALS-ENABLE beamlines are supported in part by U.S. Public Health Service award P30 GM124169. The Advanced Light Source is a Department of Energy Office of Science User Facility under Contract No. DE-AC02-05CH11231. The funders had no role in study design, data collection and analysis, decision to publish, or preparation of the manuscript.

## Author contributions

B.Z. and L.H. conceived, designed, and conducted experiments, analyzed data, contributed materials and analytic tools, and drafted the paper. S.K., N.N., C.H.L., X.S., and G.M.T. conceived and designed experiments, analyzed data, and contributed materials and analytic tools. T.S.D. and B.V.V.P. conceived and designed experiments, analyzed the data, and drafted the paper. All authors reviewed, critiqued, and provided comments on the manuscript.

## Additional information

**Supplementary information** Supplementary Table 1.

**Correspondence** and requests for materials should be addressed to Terence S. Dermody or B. V. Venkataram Prasad.

